# Interspecies interactions mediated by conductive minerals in the sediments of the ferruginous Lake La Cruz, Spain

**DOI:** 10.1101/366542

**Authors:** Amelia-Elena Rotaru, Nicole R. Posth, Carolin R. Löscher, Maria R. Miracle, Eduardo Vicente, Raymond P. Cox, Jennifer Thompson, Simon W. Poulton, Bo Thamdrup

## Abstract

Lake La Cruz is considered a biogeochemical analogue to early Earth marine environments because its water column is depleted in sulfate, but rich in methane and iron, similar to conditions envisaged for much of the Precambrian. In this early Earth analogue environment, we show that conductive particles establish a tight metabolic coupling between electroactive microbial clades. We propose that mineral-based syntrophy is of potential relevance for the evolution of Earth’s earliest complex life forms. We show that the anoxic sediment of Lake La Cruz, which is rich in biogeochemically ‘reactive’ iron minerals, harbors known electroactive species such as *Geobacter* and *Methanothrix,* in addition to other groups which have not been previously associated with an electroactive lifestyle. Slurry incubations on various substrates in the presence of conductive particles showed significant methanogenic activity, whereas incubations with non-conductive glass beads resulted in low methanogenic rates similar to slurries without added particles. In the absence of conductive particles, all tested substrates were metabolized to acetate, which accumulated to ∼10 mM. Similar to a previous study on iron-rich Baltic Sea sediments, we observed that conductive mineral additions to La Cruz slurries enabled acetate oxidation, thus preventing acetate accumulation. Acetate oxidation coupled to high methanogenic activity was only maintained in successive mud-free enrichments when these were amended with conductive minerals. In serial mud-free transfers, conductive particles conserved a consortium of *Youngiibacter-Methanothrix,* whereas *Youngiibacter* spp. died off in the absence of conductive particles. In contrast, mud-free enrichments without conductive particles ceased any metabolic activity during the second transfers. Syntrophic consortia from this early Earth analogue environment only survived in the presence of conductive particles. Mineral-mediated syntrophy could be one of the earliest evolutionary interspecies associations. Conductive minerals might have fueled metabolic exchange between cells via intercellular electron transfer prompting tight cell-to-cell associations and possibly eukaryogenesis.

## Introduction

It has been proposed that eukaryotic life arose from syntrophic interactions between Deltaproteobacteria and methanogenic archaea (López-García and Moreira, 1999; Martin and Russell, 2003; Moreira and Lopez-Garcia, 1998) in the anoxic and ferruginous (Fe-rich) early Archaean ocean (Crowe et al., 2008). Similar conditions can be found today in the anoxic deeper waters of some lakes (Crowe et al., 2008; Bura-Nakic et al., 2009; Llirós et al., 2015), including Lake La Cruz, Spain (Camacho et al., 2017; Walter et al., 2014). Most studies of these environments have focused on the phototrophic and methanotrophic communities in the water column, yet little attention has been given to either the methanogenic community buried in the sediments or the possible impact of iron-minerals on their physiology. Only recently researchers investigated the methanogenic community from Lake Matano, Indonesia which displayed high methanogenic rates when spiked with the iron-oxide, goethite (Bray et al.. 2017), however the possibility of a mineral-mediated syntrophic interaction was not assessed.

Generally, syntrophic associations are carried out indirectly, in which case electron transfer between partners is assisted by diffusible chemicals (H_2_, formate, shuttles). These classical syntrophic interactions require two partners, a bacterium capable of oxidation of complex organics to reduced compounds (i.e. H_2_), which are then retrieved by a methanogenic archaeon, which reduces CO_2_ to methane (Shrestha and Rotaru, 2014). Recent studies have shown that, sometimes, interspecies electron transfer does not require a diffusible chemical carrier. In the absence of a diffusible electron carrier, interspecies electron transfer could occur via conductive particles (magnetite, chars, pyrite) (Chen et al., 2014;

Kato and Igarashi, 2018; Liu et al., 2012, 2015; Wang et al., 2018) or directly by forging electric connections via a self-assembled extracellular network of conductive pili and c-type cytochromes between the two syntrophic partners (Rotaru et al., 2014b, 2014a; Summers et al., 2010), the latter being known as direct interspecies electron transfer (DIET). DIET was shown to be accelerated by conductive materials possibly because cells save energy by pausing the production of their own conductive extracellular network (Liu et al., 2015; Wang et al., 2018). Consequently, mineral-mediated syntrophy is energetically more favorable than the usual syntrophic associations.

It has been proposed that Fe-minerals such as pyrite helped nucleate the membranes of the earliest cells (Russell et al., 1994; Wächtershäuser, 1988a). Many membrane bound proteins involved in electron transfer through the membranes of present day cells contain FeS centers (i.e. ferrodoxins). It is therefore likely that some of the earliest FeS proteins might have played a role in electron transfer between cells.

It has been speculated that conductive-minerals also mediate the interaction between protocells with leaky cell walls and their environment, such as were probably present in the mineral-rich Archaean ocean (Lane and Martin, 2012). Interactions between cells with different metabolisms is thought to be at the origin of eukaryogenesis, as such cells compartmentalized the functions within the eukaryotic cell (López-García and Moreira, 1999; Martin and Russell, 2003; Moreira and Lopez-Garcia, 1998). In the present study, we have investigated the conductive iron-mineral dependency of interspecies interactions between bacteria and methanogens from the sediments of the Fe-rich, stratified Lake La Cruz. Specifically, we were interested in whether reactive Fe minerals would support conductive-mineral mediated interspecies interactions. As the biogeochemical setting of the lake makes it a prime early analogue (Walter et al., 2014, Camacho et al., 2017), we also discuss the context of todays mineral-mediated syntrophy as a relic of ancestral associations.

## Materials and methods

### Sampling and incubations

During an expedition at Lake La Cruz in central Spain (Fig. 1) in September 2014, we sampled the lake water and sediment. Lake La Cruz is a permanently stratified, meromictic, doline lake located in a karst region in the Iberian Mountain Range. The lake is circular with a diameter of 122 m. At the time of sampling, the maximum depth was 20 m and the chemocline was at ∼12 m depth. Water samples were pumped from sampling depths above, within, and below the chemocline at the deepest part of the lake from a boat tethered from shore to shore of the lake. The pumping apparatus was designed to withdraw water samples without contact with the atmosphere, and both the apparatus and sampling protocol have previously been described in detail (Miracle et al., 1992; Posth et al., 2017). Samples were gathered and fixed directly on the boat and stored until analysis in the lab.

**Figure 1.**
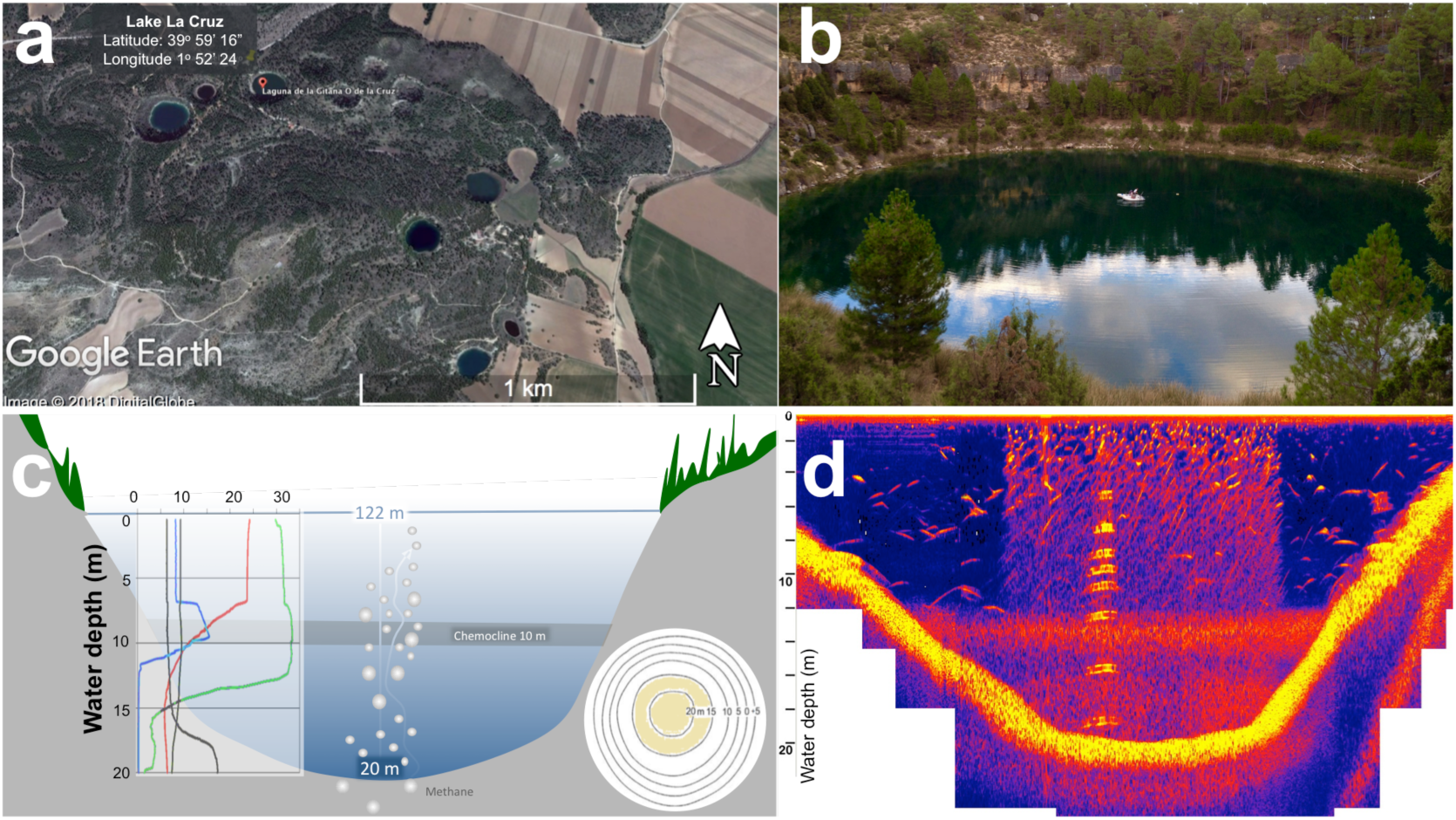
Lake La Cruz with its geophysical features. Map of the Cuenca lake area with geographical coordinates (a); and an image of the sampling site - lake La Cruz (b). Schematic representation of the lake (c) including a bathymetric map (c-round inset), and *in situ* physic-chemical characteristics of the water column (c-graphic inset). An echogram indicating the chemocline from 10-12 m, with visible gas ebullition in the central area (d).

Three sediment cores were collected from the center and deepest part of the lake (coordinates: 1° 28’17” West; 39° 59’20 North, Fig 1) using a sediment corer (Kajak sediment core, KC Denmark). The cores (50 length × 7 cm diameter) were sealed without air bubbles as they were pulled up from depth with rubber stoppers immediately inserted to avoid exposure to the atmosphere. Within 24 hours of sampling, the sediment was partitioned into depth intervals, and fixed for biogeochemical and molecular analyses inside an N2-filled inflatable glove bag, as described in detail below.

For downstream incubations, sediment from 0-15 cm depth was sampled and placed in Duran bottles secured with butyl-rubber stoppers, with a headspace of 2 bars N_2_:CO_2_ 80:20 mix. Samples were stored at 4°C until later used for incubations.

Slurries were prepared in an N_2_-filled anaerobic chamber in the laboratory. For these slurries, we used 3 mL cut-off syringes to distribute 2.5 mL of sediment into 20 mL gas-tight vials filled with 7.5 mL of medium, either modified DSM 120 or DSM 334. Modified DSM 120 medium was prepared as described previously (Rotaru et al., 2014b), but with 0.6 g/L NaCl. Three successive ten-fold dilutions of the sediment slurries led to essentially mud-free enrichments in which sediment particles could not be detected visually or by microscopy. Before inoculation, the complete medium, which lacked the substrate and (semi)conductive minerals, was dispensed anaerobically by syringe into sterile degassed vials with or without minerals prepared as below.

Two electrically conductive particle types (granular activated carbon and magnetite) were selected to be tested because they were previously confirmed to stimulate DIET in methanogenic co-cultures (Liu et al., 2012; Zheng et al., 2017). Granular activated carbon (GAC, Sigma Aldrich) had a particle size between 180 and 300 μm diameter and estimated conductivity of circa 1000 S/m (Kastening et al., 1997), and magnetite (Sigma Aldrich) with particles less than 5 diameter, and estimated electrical conductivity ranging between 0.1 and 1 S/m (Blaney, 2012; Rochelle and Schwertmann, 2003). Both materials have conductivities similar or higher than the pili that carry out extracellular electron transfer in *Geobacter sulfurreducens* (5 S/m (Adhikari et al., 2016)). 0.1 g/L of each material was weighed, added to vials, overlaid with 200 μl ultrapure water for wet sterilization, degassed for 3 minutes with N_2_:CO_2_ 80:20 mix, and autoclaved at 121°C for 25 min. Control experiments with non-conductive particle were carried out with acid-washed glass beads (less than 105 μm diameter) instead of conductive minerals. Substrates (5 mM glucose, 5 mM sodium butyrate, 10 mM sodium acetate, 10 mM ethanol) were added to media from sterile anoxic 1M stocks using aseptic and anaerobic techniques. Control experiments without electron donors were carried out in order to identify whether the organics in the sediment served as substrates for methanogenesis. All incubations were carried out at room temperature (20-23°C) in triplicate, unless otherwise noted.

Gas samples were withdrawn, stored anaerobically and then analyzed for methane on a Thermo Scientific gas chromatograph (Rotaru et al., 2018). To test for short chain volatile fatty acids (SCVFA) we used high performance liquid chromatography (HPLC) as described elsewhere (Rotaru et al., 2018).

### Biogeochemistry

For biogeochemical parameters, we took water column samples at various depth intervals and sampled the sediment obtained via the gravity corer. Geochemical parameters of relevance to this work were methane, soluble ferrous iron, and particulate reactive iron mineral species.

We will use the term reactive iron species to refer to oxalate, dithionite and HCl soluble iron oxides and sulfides (Phillips and Lovley, 1987; Poulton et al., 2004; Raiswell and Canfield, 1998).

Water column methane was sampled from the pumping apparatus through isoversinic tubing into 20 mL glass, GC vials (Supelco, Sigma-Aldrich). For each sampling depth, triplicate 5 mL samples were added to GC vials pre-doped with 10 mL 2 N NaOH to retain CO_2_ in the liquid phase. The vials were sealed with butyl-viton rubber stoppers, and stored upside down in the dark at 4°C until analysis.

Sediment methane concentrations were determined from sediment slices extracted every 2 cm in an anoxic glove bag filled with N_2_ gas.

Our measurements of available electron acceptors at the sediment boundary layer corroborated previous investigations during summer months at this lake (Camacho et al., 2017; Miracle et al., 1992; Walter et al., 2014) and showed a depletion of sulfate and Fe^3+^ (<10 μM sulfate, <1μM Fe^3+^) and no detection of oxygen and nitrate. Thus sediments mainly rely on methanogenesis for decomposition of organic matter below the water-sediment boundary, similar to previous observations on this lake during summer months (Miracle et al., 1992; Walter et al., 2014). For sedimentary methane determination, sliced sediment was filled into glass GC vials, to which 1 M (2.5%) NaOH was added in order to stop any additional microbial activity. The vials were capped with butyl-viton stoppers, crimped, and inverted until lab analysis. Sedimentary methane concentrations were determined on a Perkin Elmer GC, as previously described (Rotaru et al., 2018).

Porewater was analysed for reduced iron concentrations at ∼2 cm depth resolution after extraction using Rhizons (Rhizosphere; pore size 0.2 μm) inside a glove bag with an N_2_-atmosphere. Dissolved Fe^2+^ was determined immediately using the ferrozine assay (Lovley and Phillips, 1987; Phillips and Lovley, 1987; Stookey, 1970; Viollier et al., 2000).

To determine iron mineral speciation, sediment was subsampled from each 2 cm-depth interval and stored at -20°C. Reactive iron species (dithionite and HCl soluble iron species) (Phillips and Lovley, 1987; Poulton et al., 2004; Raiswell and Canfield, 1998) were identified from freeze-dried samples stored at -20°C by applying a modified sequential iron extraction procedure (Poulton and Canfield, 2005). In the first step, a room temperature 0.5 N HCl extraction was applied to dissolve poorly crystalline iron oxides such as ferrihydrite, surface absorbed Fe^2+^, iron carbonate minerals such as siderite, and acid volatile iron monosulfides (Zegeye et al., 2012). Subsequently, a pH 4.8 sodium dithionite extraction was employed to dissolve crystalline ferric oxide minerals such as goethite and hematite, followed by an oxalate extraction to dissolve magnetite (Poulton and Canfield, 2005). The total concentration of iron dissolved in each operationally defined extraction phase was determined by flame atomic absorption spectroscopy (AAS). For the 0.5 N HCl extraction, dissolved Fe^2+^ was also measured immediately via the ferrozine assay (Lovley and Phillips, 1987; Phillips and Lovley, 1987; Stookey, 1970; Viollier et al., 2000). Extraction of this Fe^2+^ from the total Fe determined for this extraction by AAS gave the Fe^3+^ concentration associated with poorly crystalline iron oxides such as ferrihydrite. Iron sulfide phases were determined via a two-step sequential extraction procedure (Canfield et al., 1986). Acid volatile sulfide minerals (FeS) were determined by extraction with hot 6 N HCl under N2, with the released sulfide trapped as Ag_2_S. Pyrite (FeS_2_) was then determined after addition of chromous chloride, with the sulfide again trapped separately as Ag_2_S. After filtration, the concentrations of Fe in FeS and FeS_2_ were determined stoichiometrically. The concentration of Fe present as FeS was subtracted from the Fe^2+^ concentration determined by the 0.5 N HCl extractions, to give surface reduced and carbonate-associated Fe^2+^. Replicate extractions gave a RSD of <5% for all phases.

### Scanning electron microscopy

Samples from the water column were preserved in 4% formalin, filtered on Nucleopore carbonate filters, with a pore size of 0.2 μm, and dehydrated in 20 minute steps with ethyl alcohol (30%, 50%, 70%, 90% and two times 100%). Then samples were critical point dried prior to palladium/gold sputter coating and visualization on a Hitachi S-4800 FE scanning electron microscope ran at an electron beam acceleration voltage of 10kV.

### Molecular analyses

For molecular analyses we sampled 2 mL of sediment at 2 cm depth resolution using cut-off syringes inside a N_2_-filled glove bag. Sediment was pooled together every 4 cm and fixed with MoBio RNAlater 1:1 v/v (Rotaru et al., 2018). Prior to DNA extractions, most of the RNAlater was removed by centrifugation. For DNA extraction we used the top 16 cm of sediment from triplicate cores. Extractions were carried independently for each core with the MoBio RNA Soil kit coupled to the MoBio complementary DNA Soil kit, following the manufacturer’s protocol. DNA was quantified using a Nano Drop prior to downstream applications. The DNA extracted from each core was amplified with the following primer pair S-D-Arch-0519-a-S-15/ S-D-Bact-0785-b-A-18, which according to Klindworth et al. (2013), was the best for MiSeq amplicon sequencing, targeting more than 89% of Bacteria and more than 88% of Archaea. PCR amplification and indexing (using Nextera XT index kit, Illumina) of the PCR products for the triplicate samples was conducted following the Illumina 16S rRNA gene amplicon sequencing protocol (Illumina, USA). The DNA samples were then sequenced using ×300 PE MiSeq sequencing approach at Macrogen (www.macrogen.com), using Illumina’s protocol. The sequences generated circa 1 million reads for each core, which were imported into CD-HIT-OTU to remove noisy data and clustered into OTUs, using a 97% species cutoff. For taxonomy and diversity analyses, clean and clustered OTUs were analyzed using QIIME (Caporaso et al., 2010), against the Ribosomal Database Project database version 11. Alpha rarefaction analyses showed sufficient coverage of the diversity in all three sediment cores.

DNA extractions from mud-free incubations were performed using the MasterPure DNA purification kit as previously described (Rotaru et al., 2014b). Amplification of bacterial (27F, 5'-AGAGTTTGATCMTGGCTCAG and 1492R, 5'-TACCTTGTTACGACTT) and archaeal (344F - 5’ -ACGGGGYGCAGCAGGCGCGA-3’ and 1059R – 5’- GCCATGCACCWCCTCT-3’) 16S rDNA sequences, library preparation, and 16S rRNA gene sequencing, was performed as previously described (Rotaru et al., 2018). Maximum likelihood phylogenetic trees were constructed using Geneious (Kearse et al., 2012). Sequence files can be found at NCBI under Bioproject ID: PRJNA (in the process of submission).

## Results and discussion

Our hypothesis was that the iron-rich Lake La Cruz would be the breeding ground for conductive, mineral-based syntrophy (Rotaru et al., 2018). We discovered that microorganisms enriched from Lake La Cruz carried out syntrophic degradation strictly dependent on conductive mineral additions and were unable to carry unaided DIET associations.

### Geochemistry

We expected to find a niche for DIET/conductive-particle mediated IET in this Fe-rich methanogenic lake resembling the ocean in the Precambrian. La Cruz sediments displayed high methane concentrations in the top 15 cm, along with a significant proportion of reactive iron species ∼70% of the total Fe content) (Thompson 2018), which is very high relative to normal non-ferruginous aquatic environments (Poulton and Raiswell, 2002; Raiswell and Canfield, 1998). During this sampling campaign, the sediments were overlain by ∼10 m of anoxic water (Fig. 1). During summer months, the lake is known to persistently have a 4-5 m monimolimnion zone above the sediment, which is rich in Fe^2+^ (Vincente and Miracle, 1988). In our study we also noticed a strong methane supersaturation near the bottom, where the methane concentration reached 4 mM, similar to concentrations in the surface sediment (Fig. 2). Gas ebullition from the deep water table during sampling and oftentimes gas bubbles, mainly consisting of methane and carbon dioxide (Camacho et al.. 2017), percolated through the surface of the lake from the middle, as documented by a ecogram of the lake (Fig. 1d). Previous studies suggested that the sediment is the source of water-column methane (Oswald et al., 2016). Indeed we observed that sediment methane concentrations were highest in the top centimeters of the sediment (Fig. 2). Methane concentrations were also high in the water column (17-20 m), indicating methanogenesis occurs in the bottom waters as well as the top layers of the sediment (Fig. 2).

**Figure 2.**
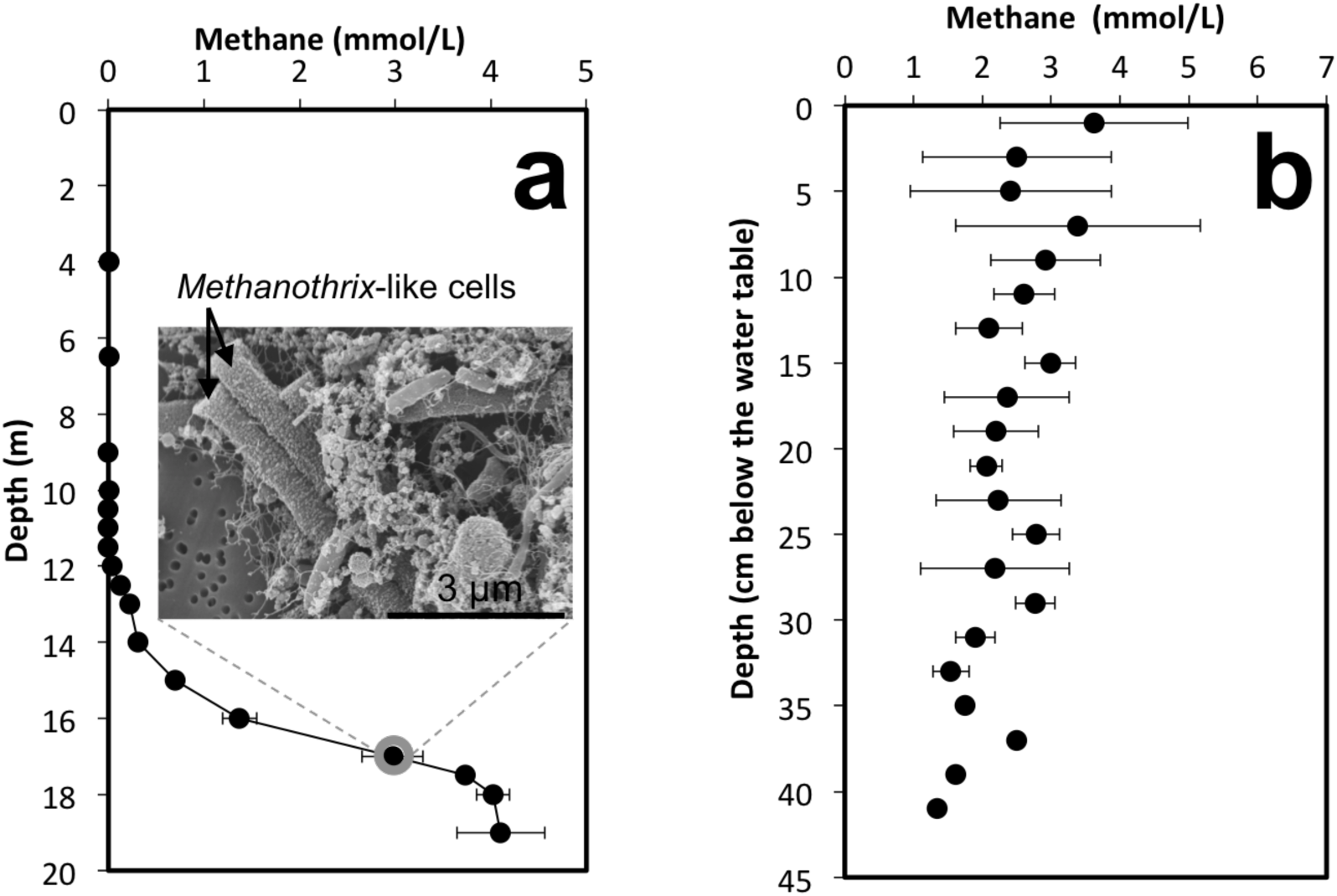
Methane profiling through the water column and sediment of lake La Cruz. (a) In the water column of Lake La Cruz, the highest methane concentrations were below 17-m depth where *Methanothrix*-like cells (inset) could be observed by scanning electron microscopy. (b) Sediment cores showed very high methane concentrations especially in the top 15-cm, indicating that methane also has sedimentary origin. The water column average values are for triplicate samples taken at each specific water column depth; while values for sediments are from triplicate cores sampled every 2-cm.

Similar to previous studies (Camacho et al., 2017, Oswald et al., 2016), dissolved Fe^2+^ builds up below the chemocline to reach concentrations of ∼250 μM above the sediment-water interface. In the sediment, dissolved Fe^2+^ concentrations continue to increase (Thompsen 2018), reaching a peak of >1000 μM at a depth of ∼22 cm. These high dissolved Fe^2+^ concentrations in the water column and sediment porewaters are similar to those found in other iron-rich lakes (Bura-Nakic et al., 2009; Crowe et al., 2011; Nordi et al., 2013; Vincente and Miracle, 1988). The La Cruz sediments were high in TOC (average = 6.68 ± 2.0 wt %), and carbonate minerals (average = 9.46 ± 1.3 wt% inorganic C) which diluted the total Fe-content to 1.06 ± 0.18 wt% on average (Thompson, 2018). This is significantly lower than the average global total Fe content of riverine particulates supplied to oceans and lakes (4.49 wt%; Poulton and Raiswell, 2002). Proportionally, however ‘reactive’ Fe phases (non-sulphidized Fe^2+^, Fe-oxides, Fe-sulfides) were abundant (70±8%; Thompson, 2018) relative to the total Fe content of the sediment, of which only 18±5% was sulfur bound (pyrite, other Fe-S minerals). Nevertheless, iron-oxide concentrations were rather low, with magnetite Fe accounting for less than 0.1% of the total Fe-content in this sediment. Other Fe oxide minerals accounted for ∼10% of total Fe on average. Thus, non-sulphidized particulate Fe(II) was the dominant reactive Fe pool (∼60% on average).

Some of the iron minerals (Fe-sulfides and Fe-oxides) found in the sediments of lake La Cruz are electrically conductive (Fig. 3), of which magnetite and iron sulfides were documented to facilitate mineral mediated syntrophy (Kato and Igarashi, 2018; Liu et al., 2012, 2015; Rotaru et al., 2018; Zheng et al., 2017). Fe-sulfides, like pyrite were also shown to aid long-range extracellular electron transfer from cells (Kondo et al., 2015) or enzymes (Mahadevan and Fernando, 2018) to electrodes. Moreover La Cruz sediments also contain coal particles (Romero-Viana et al., 2011), which are conductive (Fig. 3). Indeed it has been documented that conductive carbon materials (i.e. granular activated carbon) facilitated mineral mediated syntrophy as effectively as conductive Fe-minerals (Liu et al., 2012; Rotaru et al., 2018).

**Figure 3.**
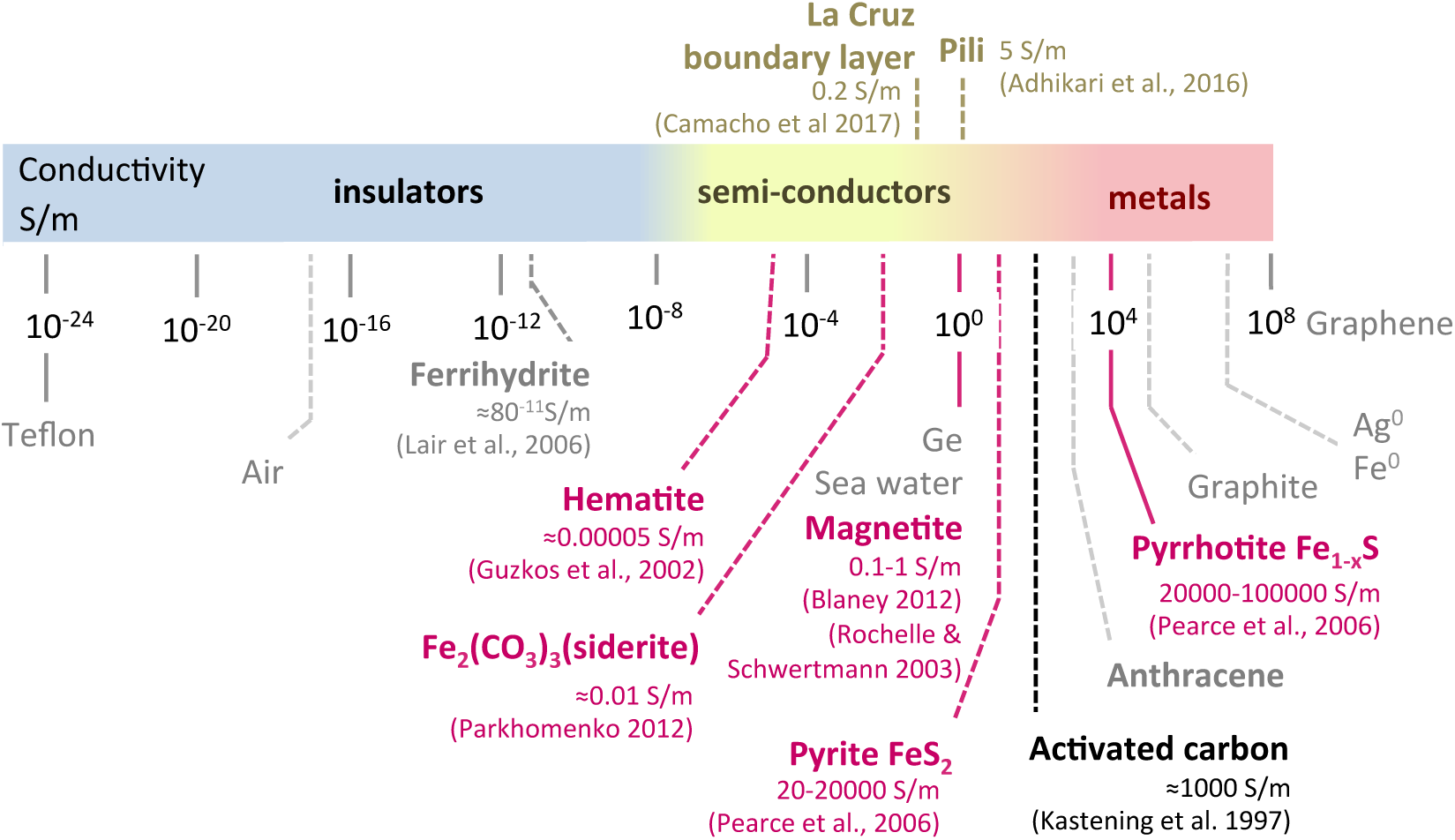
Conductivities of various Fe-oxides and Fe-sulfides, compared to that of the e-pili and the conductivities observed for different carbon particles including activated carbon used in this study. We have also listed the conductivity measured in lake La Cruz at the sediment-water boundary layer. (Adhikari et al., 2016; Blaney, 2012; Camacho et al., 2017; Guskos et al., 2002; Kastening et al., 1997; Lair et al., 2006; Parkhomenko, 1990; Pearce et al., 2006; Rochelle and Schwertmann, 2003)

### *In situ* bacterial diversity - with focus on described electrogens

We therefore anticipated that electrically conductive particles inherent to La Cruz sediments would facilitate mineral mediated interactions between electrogens and electrotrophic methanogens in this lake sediment. Indeed, our data demonstrate that the community harbours organisms affiliated to groups of electrogens including *Geobacter* (Fig. 4), and to DIET-methanogens including *Methanothrix* (Fig. 4). *Geobacter* and*Methanothrix* have previously been shown to carry out direct interspecies electron transfer in laboratory co-cultures (Rotaru et al., 2014a), and have been found to co-exist in several man-made environments, such as rice paddies (Holmes et al., 2017) and anaerobic digesters (Morita et al., 2011; Rotaru et al., 2014a). In this study we show that bacteria affiliated to known electrogens/iron-reducers like *Geobacter* (0.6% of all Bacteria), *Thiobacillus* (0.2% of all Bacteria), *Desulfobacterium* (0.4% of all Bacteria), and *Anaerolinea* (0.1% of all Bacteria) co-exist with *Methanothrix* in Lake La Cruz sediments (Fig. 4). Together, all of these putative electrogens/iron reducers were represented in Lake La Cruz sediments, summing up to circa 1% of all Bacteria. Previously, members of these four genera, *Geobacter, Thiobacillus, Desulfobacterium, Anaerolinea,* were shown to be capable of extracellular electron transfer to and/or from electrodes or metallic iron (Dinh et al., 2004; Gregory et al., 2004; Kawaichi et al., 2018; Nakasono et al., 1997; Pous et al., 2014; Rotaru et al., 2015), as well as iron-minerals (Bosch et al., 2012; Kawaichi et al., 2013; Lovley et al., 1993; Rotaru et al., 2015). The first two, *Geobacter* and *Thiobacillus* can also interact by DIET with other cells (Kato et al., 2012; Rotaru et al., 2014b, 2014a; Summers et al., 2010), and this interaction has been shown to be expedited in the presence of conductive particles (Chen et al., 2014; Kato et al., 2012; Liu et al., 2012, 2015; Rotaru et al., 2014b; Zheng et al., 2017). It is therefore possible that all of these electrogenic species compete for the electron uptake of electrotrophic methanogens.

**Figure 4.**
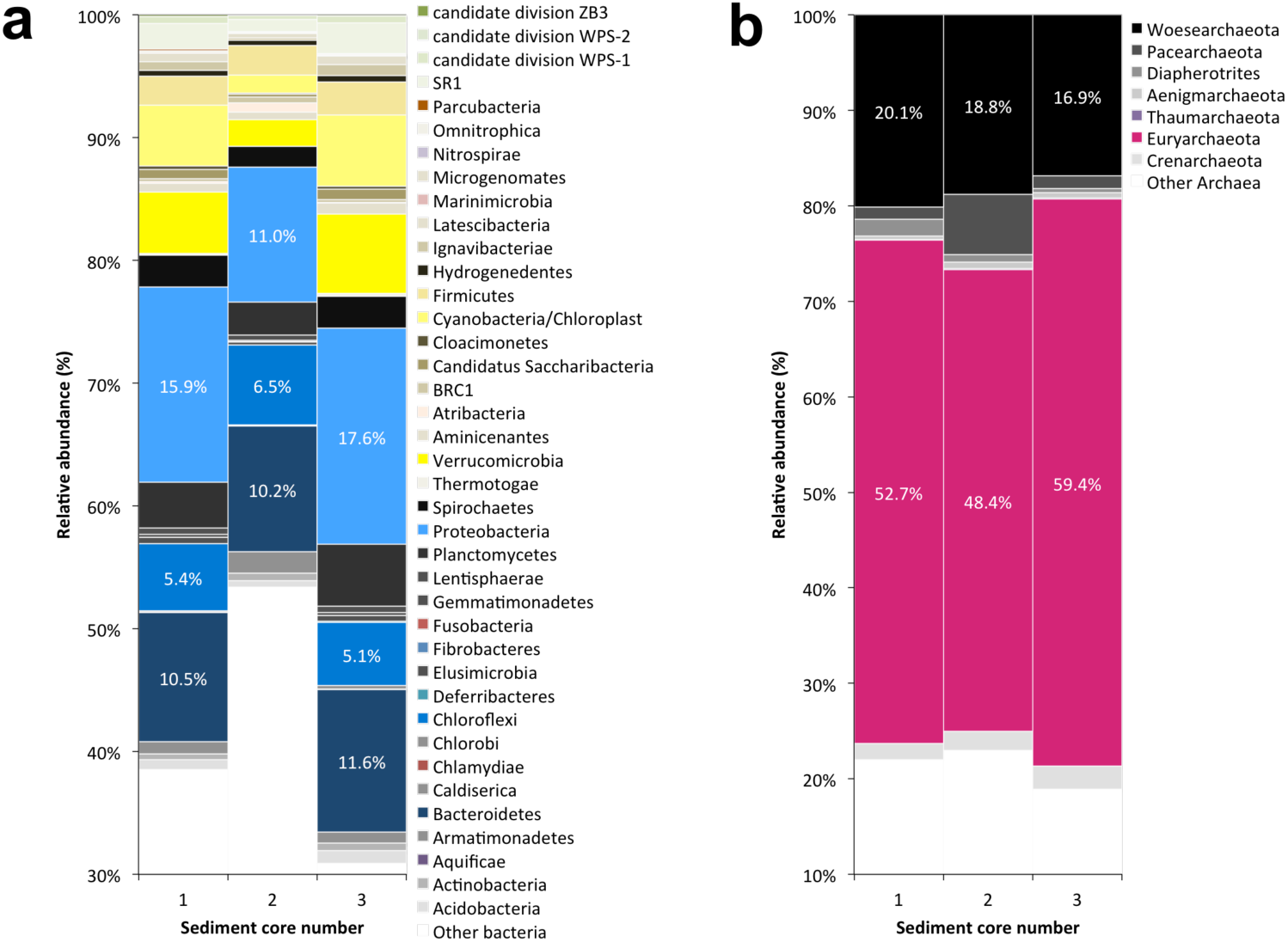
Relative, phylum-level composition of (a) Bacteria and (b) Archaea harboring the top 16-cm of three sediment cores from lake LaCruz, as determined by 16S rRNA gene amplicon sequencing.

However, one of the most abundant genera in these sediments was *Smithella* (2.6% of all Bacteria), which were also potentially electroactive and may carry DIET interactions with *Methanothrix* in an alkane-degrading consortium (Embree et al., 2014). Therefore, *Smithella* may establish a DIET-association with *Methanothrix* from Lake La Cruz sediments (see Archaea community below). Members of *Verrucomicrobia* were also very well represented (circa 4.6% of all Bacteria) similar to what has been observed for 90% of several lake sediments (He et al., 2017). *Verrucomicrobia* were recently proposed to carry extracellular electron transfer due to their genetic make-up, which comprises the appropriate porin systems and membrane-associated c-type cytochromes (He et al., 2017). It is also possible that *Verrucomicrobia* play a role in mineral mediated interspecies interactions. Nevertheless, *Verrucomicrobia* has never been shown to have the ability to interact syntrophically or to carry out extracellular electron transfer in laboratory cultures, and thus this predicted physiology requires future investigation. Some of the most abundant phyla were Bacteroidetes (10.8% of the Bacteria), and Firmicutes (2.5% of all Bacteria) (Fig. 4).

### *In situ* archaeal diversity

*Euryarchaeaota* accounted for more than half of the *Archaea* represented through amplicon sequencing (Fig. 4). Here, we show that in the sediments of Lake La Cruz, *Methanothrix* co-existed with electrogens *(Geobacter, Thiobacillus, Desulfobacterium,* and *Smithella).* Besides the acetoclastic/DIET-associated *Methanothrix* (3.7% of all *Archaea),* we identified canonical hydrogenotrophic-methanogens belonging to *Methanoregula* (2.5% of all *Archaea),* and very low numbers of*Methanobacterium* (0.2% of all *Archaea).* The most abundant *Archaea* were the deep-branching *Methanomassilicoccus* (40.6% of all *Archaea).* The role of*Methanomassillicoccus* in sedimentary methanogenesis is not well understood since their only cultivated species-representative, *M. luminyiensis,* is a human-gut isolate strictly capable of H_2_-dependent methylotrophic methanogenesis, but incapable of CO_2_-reductive methanogenesis or acetoclastic methanogenesis (Dridi et al., 2012a). Besides their documented presence in the human gut (Adam et al., 2017; Dridi et al., 2012b), *Methanomassilicoccus* species have also been found in the guts of insects (Paul et al., 2012) and animals (Raymann et al., 2017; Salgado-Flores et al., 2018; Söllinger et al., 2016), anaerobic digesters (Chojnacka and B, 2015; Kuroda and Hatamoto, 2015), hydrothermal springs (Coman et al., 2013; Merkel et al., 2015, 2016), wetlands (Söllinger et al., 2016), subsurface aquifers and soils (Kadnikov et al., 2017; Rout et al., 2015), and riverine and marine sediments (Guo et al., 2018; Nunoura et al., 2016; Rotaru et al., 2018; Vigneron et al., 2016). *Methanomassiliicoccus* was also one of the most abundant genera of methanogens, not only in the iron-rich sediments of Lake La Cruz, but also in Baltic Sea sediments that are potential niches for conductive particle-mediated syntrophy (Rotaru et al., 2018). It is possible that *Methanomassilicoccus* is involved in electroactive interactions via minerals, especially taking into account that this group was recently associated with electroactive communities abundant on electrodes from bioelectrochemical systems set up with inoculums from soils (Ahn et al., 2014) and anaerobic digester sludge (Park et al., 2018).

Among the methanogens detected in La Cruz sediments, only species of *Methanothrix* have been previously shown to establish DIET-associations with *Geobacter* species (Holmes et al., 2017; Morita et al., 2011; Rotaru et al., 2014a; Wang et al., 2016). *Methanothrix* was earlier suggested to carry out DIET with *Smithella* (Embree et al., 2014), but the latter has never been shown to be capable of mineral-mediated or direct electron transfer. In a previous study, we have shown that a *Methanothrix-species* from the Baltic did not establish a mineral-mediated interaction with Baltic-*Geobacter*, but were instead competitively excluded by a *Methanosarcina-Geobacter* consortium, which carried a mineral-mediated syntrophic association (Rotaru et al., 2018). However, although *Methanosarcina* is a very effective DIET partner (Rotaru et al., 2014b, 2015) and mineral-syntrophy partner (Chen et al., 2014; Liu et al., 2012; Rotaru et al., 2018; Wang et al., 2018) they were poorly represented in La Cruz sediments (Fig. 4).

### High methanogenic activity could only be maintained by conductive particles

In order to determine the effect of conductive particles on the Lake La Cruz methanogenic community, we compared incubations with or without additional conductive particles. These incubations showed that the methanogenic community was strictly dependent on the addition of conductive particles and independent of the type of substrate, conductive particle, or freshwater medium tested (Fig. 5). Incubations with conductive particles showed 2-4 fold increases in methanogenic rates (0.2-0.7 mM/day, depending on substrate) over incubations with non-conductive glass beads or without particle-amendment (0.09 to 0.18 mM/day, depending on the substrate). Moreover, high methanogenic activity was maintained in subsequent incubations *only if* conductive particles were added (Fig. 5). Cultures without conductive particles could not sustain methanogenesis for more than one subsequent transfer. This indicates a strict dependency of the enriched methanogenic community on conductive particles.

**Figure 5.**
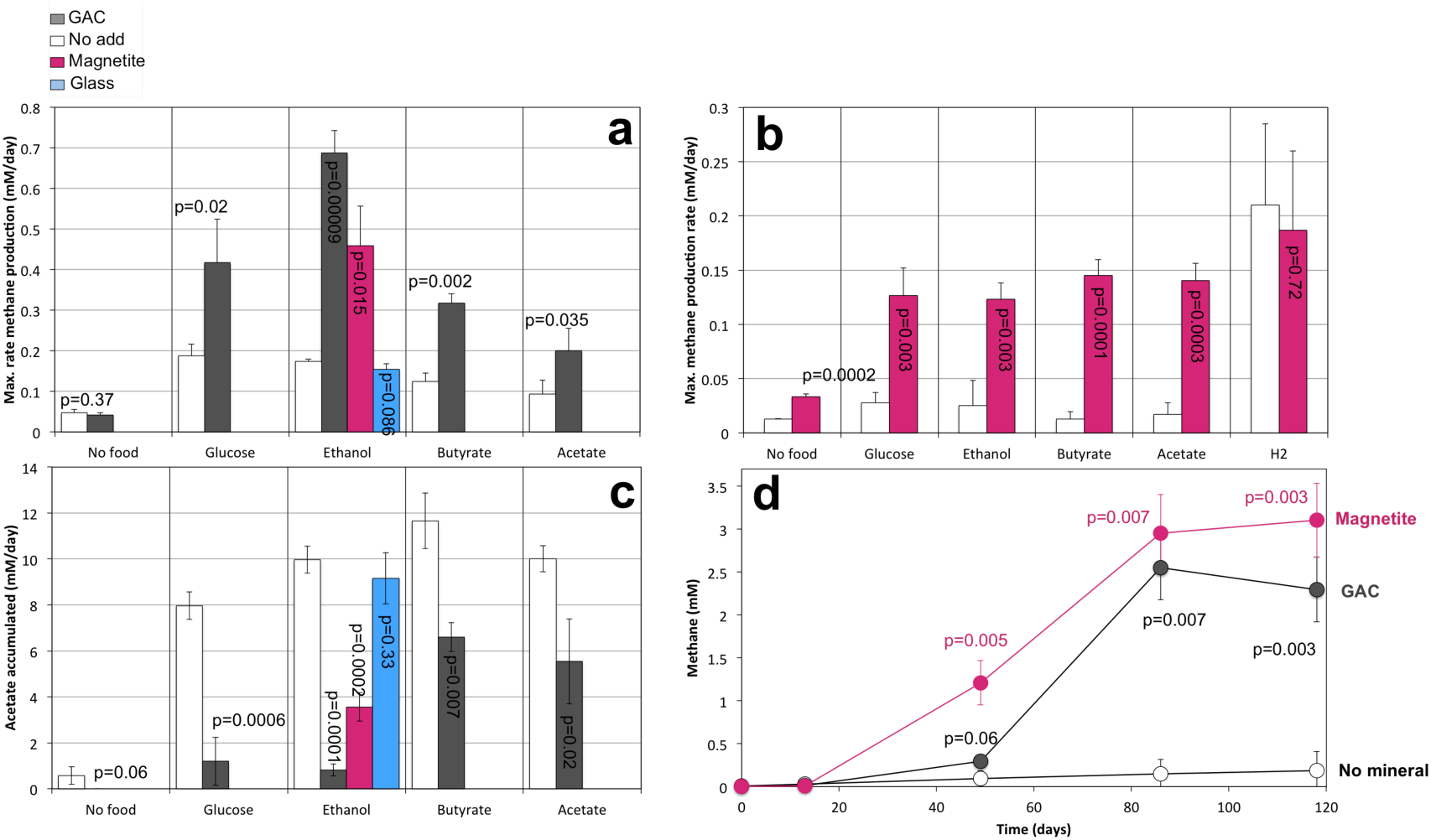
Methanogenesis on different substrates in incubations from lake La Cruz. Methane production in initial slurry incubations provided with different substrates was stimulated by conductive particles (GAC or magnetite) independent of the media used (a) modified DSMZ 120 or (b) DSMZ 334. (c) Acetate accumulated in incubations without conductive particles, but was significantly lower at the addition of conductive particles. (d) For example, a third transfer free of sediment showed that methanogenesis and acetate consumption were strictly dependent of the presence of conductive particles (colored symbols), and ceased if conductive particles were not added consistently for subsequent transfers (white symbols).

We observed that all tested substrates were transiently converted to acetate, which was converted quickly to methane in the presence of conductive particles, whereas acetate accumulated in the absence of conductive particles (Fig. 5). This is likely due to higher rates of acetate oxidation prompted by the addition of conductive particles, similar to previous observations of Bothnian Bay sediments where syntrophic acetate oxidation (SAO) relied on conductive minerals (Rotaru et al., 2018).

We determined which organisms were enriched on acetate with or without conductive particles. For this we compared the acetate fed communities exposed to two types of conductive particles (GAC and magnetite) to a community exposed to no conductive particles. We determined that *Youngiibacter* and *Methanothrix* methanogens dominated the enrichments amended with both types of conductive particle (Fig. 6). On the other hand, in controls without conductive particles, after only one single transfer *Youngiibacter* could not be detected. In the absence of conductive particles methane production only occurred slowly for one transfer and in this case *Methanothrix* co-existed with *Clostridium* (Fig. 6).

**Figure 6.**
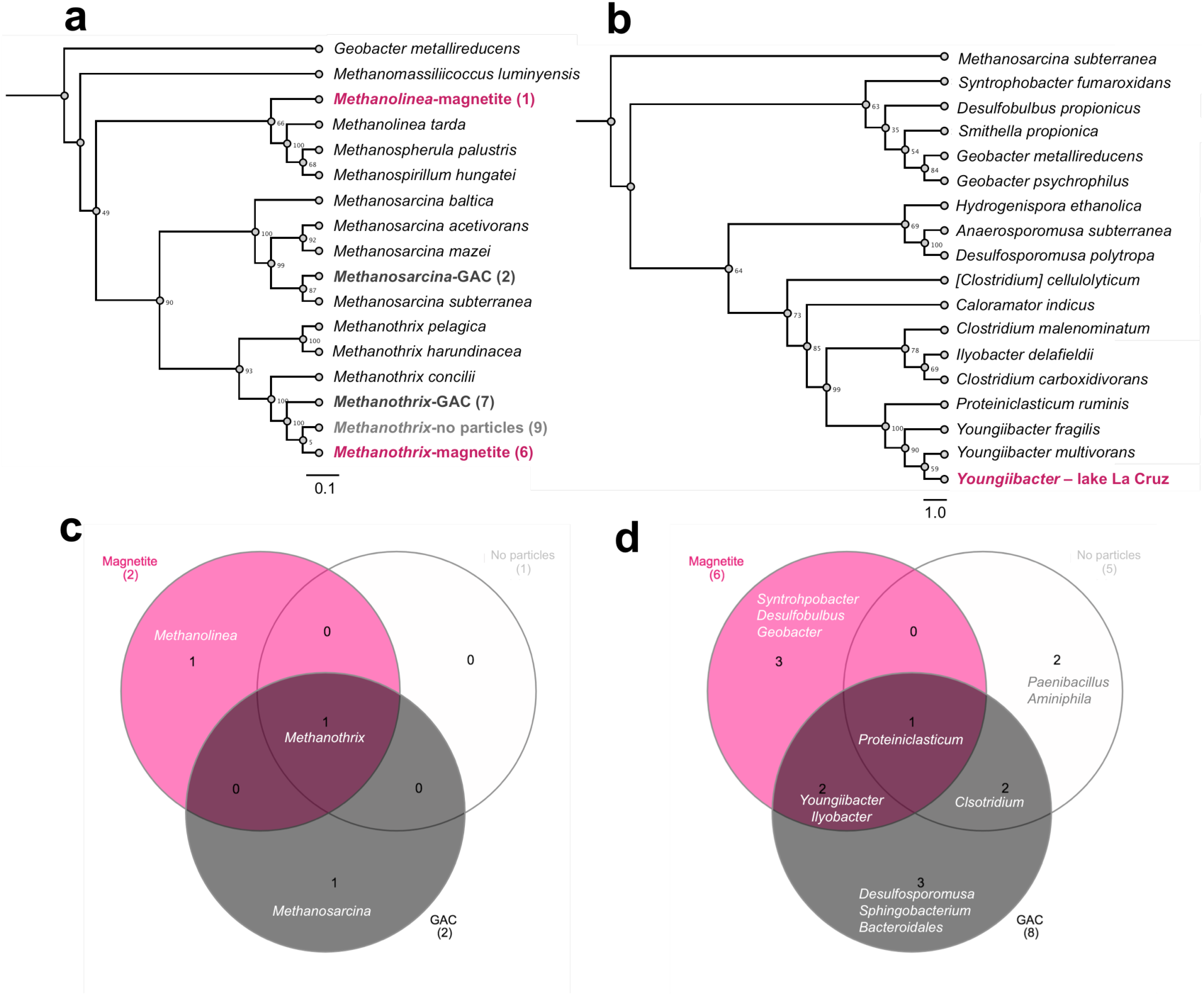
Maximum likelihood phylogenetic trees (a, b) and venn diagrams with the relative distribution of 16S rRNA-gene sequences for Archaea (a, c) and Bacteria (b, d) in La Cruz incubations with or without conductive particles. Archaeal (a) and Bacterial (b) 16S rRNA genes were retrieved from third mud-free transfer of acetate-incubations with magnetite (pink), and GAC (black-bold) or from a first mud-free transfer without conductive particles (light gray-white). (c) The only Archaeal 16S phylotype encountered in all incubations independent of treatment was *Methanothrix-related.* (d) The most abundant Bacterial phylotype encountered only in conductive particle-amended incubations was *Youngiibacter*-related.

*Youngiibacter* was only found in enrichments with conductive particles and its presence could be associated with rapid acetate consumption coupled to methane production (Fig 5). We therefore anticipate that *Youngiibacter* plays a role in conductive-particle mediated syntrophy. Nevertheless, until now little is known about this group of Firmicutes, and only recently two species of *Youngiibacter* have been described (Lawson et al., 2014; Tanaka et al., 1991), of which one is associated with fermentation of organics on coal surfaces during coal conversion to natural gas (Lawson et al., 2014). Coal, similar to activated carbon, is electrically conductive (Duba, 1977). Moreover, *Methanothrix* have been also found associated with coal conversion to natural gas (Beckmann et al., 2011; Lawson et al., 2014). It is therefore possible that *Youngiibacter* and *Methanothrix* play a role in conductive particle-mediated syntrophy in coal beds, and as well in Lake La Cruz sediments. However, a syntrophic association between *Youngiibacter* and *Methanothrix* has not been described before. We suggest that *Youngiibacter* released electrons from substrate/acetate oxidation onto conductive minerals that are then used as a source of electrons for *Methanothrix* in order to reduce CO_2_ to methane. It is possible that *Youngiibacter* releases electrons extracellularly using a similar mechanism to that described for *Geobacter* namely a network of outer membrane c-type cytochromes and pili (Shrestha et al., 2013). During DIET, OMCs were not as necessary for a donor *Geobacter* strain to carry substrate oxidation coupled with extracellular electron transfer and respiration, since OMCs could be completely replaced by the conductive iron oxide, magnetite (Liu et al., 2015). Instead, when it plays the role of electron donor Geobacter seems to necessitate e-pili for long range electron transfer to partner cells, as exemplified in a recent study (Ueki et al., 2018). In agreement with previous observations in *Geobacter* (Ueki et al., 2018), *Youngiibacter* might employ type IV pili for EET to partner *Methanothrix. Youngiibacter’s* type IV pili gene sequence (T472_0202395) differs significantly from that of *Geobacter* species, yet it has a high content of aromatic aminoacids (10.3%) which could give this organism an advantage to carry EET (Walker et al., 2018). It is possible that conductive particles ornate the pili of *Youngiibacter* in a similar way to how they do for *Geobacter* (Liu et al., 2015; Wang et al., 2018) facilitating electron transfer to syntrophic partner methanogens.

### Conductive-particle mediated syntrophy

Syntrophy mediated by conductive particles could occur in three different ways (Fig. 7). A first mode of action includes electrogens with limited expression of surface cytochromes whose role would be replaced with that of conductive minerals (pyrite, pyrrhottite, magnetite, goethite) found in sediments (Fig. 7a). Molecular and microscopic evidence for this type of association has been brought by studies in laboratory *Geobacter* co-cultures provided with magnetite (Liu et al., 2015). A second possibility is that cells plug into macro-sized conductive rocks (i.e. iron/manganese-nodules) with one cell releasing electrons onto the rock and the other receiving electrons (Fig. 7b). Evidence for such interactions was previously obtained in laboratory co-cultures with macro-sized conductive chars. In this case, using SEM, it was shown that the electrogen/*Geobacter* cells did not require direct contact to the electrotroph/*Methanosarcina* yet the conductive surface facilitated the syntrophic association (Chen et al., 2014; Liu et al., 2012). The third possibility (Fig. 7c), is that membrane-bound proteins facilitate the precipitation of Fe^2+^-ions, i.e., with thiol groups (Milner-White and Russell, 2005) to form a conductive surface-conduit surrounding the cell. Extracellular electron transfer between such mineral-coated cells has been proposed (Kato et al., 2012), but has not been confirmed. However, this could be a possibility for microbes without an extracellular apparatus for electron transfer to partner cells.

**Figure 7.**
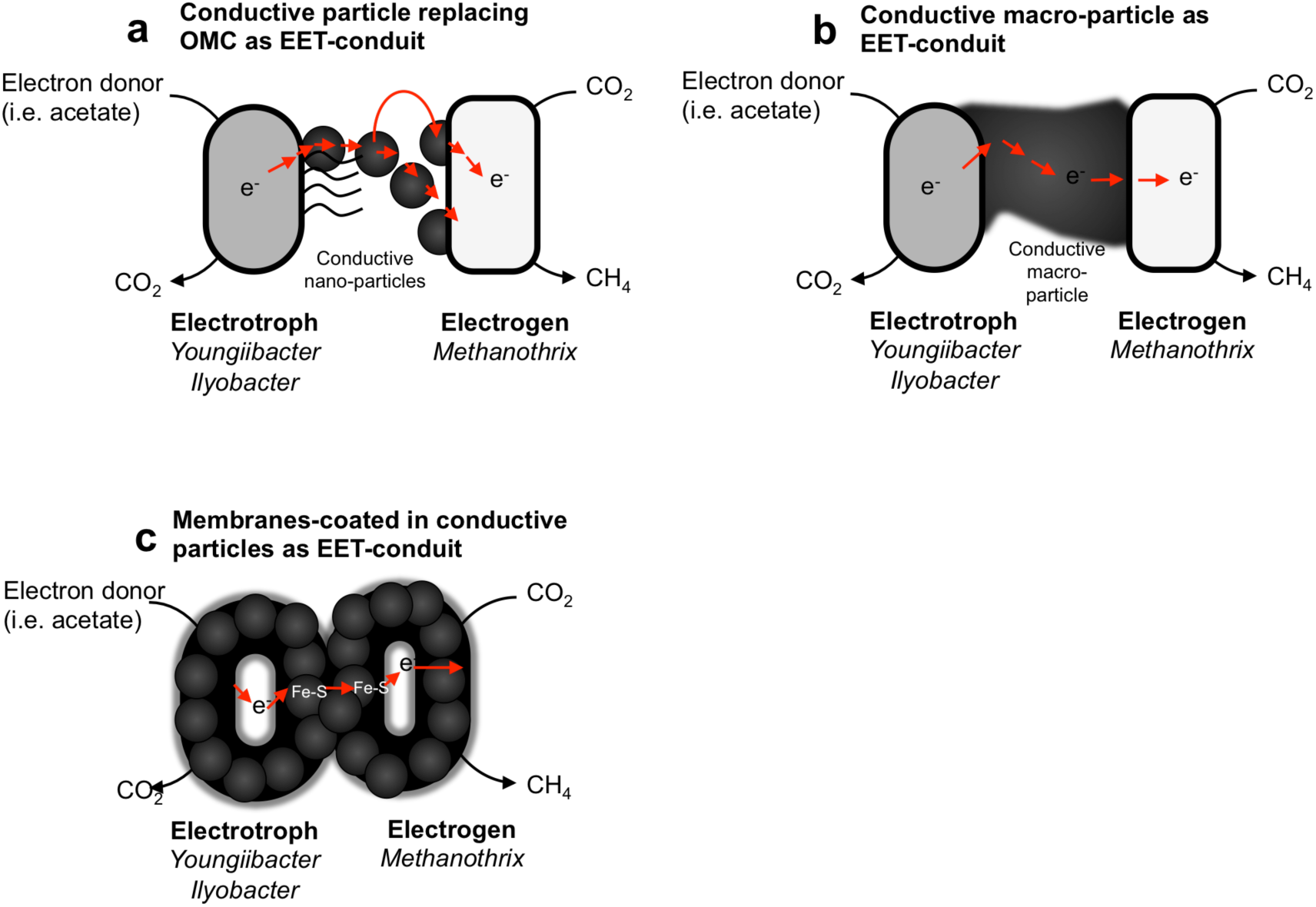
Proposed model interspecies interactions in La Cruz sediments facilitated by conductive particles. (a) Syntrophy mediated by a conductive nano-particles replacing outer membrane cytochromes (OMCs). Nevertheless, pili involved in EET are still available. (b) Syntrophy mediated by a conductive macro-particle (i.e. GAC), which plays the role of both and electron plug and outlet. (c) Syntrophy mediated by a conductive-mineral coat padding the cell surface. In lake La Cruz, conductive minerals could for example result from the precipitation of Fe^2+^ as Fe-S/thiola- in the periplasmic space of cells. Cell surfaces encrusted with a metal-S coat such as Fe-S might endorse the electron-transfer between the two distinct metabolic entities even in the absence of a typical EET/DIET conduit.

It is plausible that mineral-mediated interactions preceded in evolutionary terms the interspecies electron transfer interactions based on diffusible chemicals, which require complex enzymes and cell-bound electrical conduits. Primordial protocells had not developed enzymatic machineries to maintain redox and proton gradients across cell membranes (Martin et al., 2003; Russell et al., 1990, 1994; Wächtershäuser, 1988b). It has therefore been suggested that minerals, which can uphold voltage differences, such as FeS/pyrite, might have helped nucleate the earliest membranes, playing the role of early membrane-bound catalysts, instead of electron transport chain enzymes today (Martin et al., 2003). Later, the high reactive iron content of the Archaean ocean could have promoted the formation of proteins with Fe-S centers which are required and abundant in redox proteins of methanogens (Liu et al., 2010). Here, we propose that primitive cells with leaky membranes (Lane and Martin, 2012), allowed easy electron transfer via conductive minerals permitting energy exchange between separate metabolic protocell entities. Thus conductive particles could have fostered the earliest interspecies interactions in the methanogenic and iron-rich Early Earth oceans, and possibly nurtured adaptation of interspecies associations pre-eukaryogenesis.

## Conclusion

In conclusion, we show that the sediment of an early Earth ocean analogue is the niche for syntrophic associations dependent on conductive particles. *Only if* conductive particles were provided, could syntrophic bacteria coupled to methanogens oxidize their substrates. Thus, only in incubations with conductive particles members of the genus *Youngiibacter* were identified to co-exist with *Methanothrix.* Incubations without conductive particles resulted in the disappearance of *Youngiibacter*, and one transfer later to the demise of the methanogenic community. These data indicate that conductive particles were required to aid the pairing of the metabolism of *Youngiibacter* with that of *Methanothrix,* which sustained high rates of methanogenesis in this early Earth analogue - lake La Cruz. We propose that obligate mineral-syntrophy is an ancestral interspecies interaction established before complex membrane structures and enzymes evolved to intermediate direct or indirect associations between species with distinct metabolism.

## Acknowledgements

This work is a contribution to a Danish Research Council grant 1325-00022 awarded to AER. During the writing of this manuscript, AER has been supported by three other grants: a Sapere Aude grant from Danish Research Council (4181-00203), a Novo Nordisk Foundation award and an Innovationsfonden grant (4106-00017), CRL was supported by the EU’s H2020 program (#704272, NITROX). NP thanks the Seventh Framework Programme of the European Union Marie Skłodowska-Curie Intra-European Fellowships (BioCTrack 330064) for their support. JT acknowledges support from a NERC research studentship. We would like to acknowledge lab support by Lasse 0rum-Smidt, Erik Laursen, Heidi Gran Jensen, Bente H0lbeck, and Susanne M0ller.

